# Peptide Secondary Structure Prediction using Evolutionary Information

**DOI:** 10.1101/558791

**Authors:** Harinder Singh, Sandeep Singh, Gajendra Pal Singh Raghava

## Abstract

**BACKGROUND:** In the past, large numbers of methods have been developed for predicting secondary structure of proteins. Best of author’s knowledge no method has been specifically developed for predicting secondary structure of peptides. We analyzed secondary structure of peptides and proteins; it was observed that same peptide in protein adopt different secondary structures. Considering the wide application of peptides in therapeutic market, we made attempt to develop a method called PEP2D for predicting secondary structure of peptides.

**RESULTS:** In this study, 3107 unique peptides have been used to train, test and evaluate peptide secondary structure prediction models. It was observed that regular secondary structure content (e.g., helix, beta-sheet) increased with length of peptides. Firstly, models based on various machine-learning techniques have been developed using binary profile of peptides and achieved maximum overall accuracy (Q3) 79.5%. The performance of models further improved from 79.5% to 83.5% using evolutionary information in the form of PSSM profile. We also evaluate performance of protein secondary structure prediction method PSIPRED on our dataset and achieved maximum accuracy 76.9%; particularly poor (Q3 71.4%) for small peptides having length less than 10 residues. Overall, PEP2D has better prediction of beta-sheets (Q3 74%) and coil region (Q3 87%) of peptides as compare to PSIPRED (Q3 54.4% for beta-sheet and Q3 77.9% for coil). We also measure performance of PSIPRED and PEP2D in terms of segment overlap (SOV); achieved 69.3 and 76.7 respectively.

**CONCLUSION:** Our observations indicate that there is a need of developing separate method for predicting secondary structure of peptides. It was also observed that models based on PSSM profile perform poor on small peptides in comparison to long peptides. Based on our study, we developed method for predicting secondary structure of peptides. In order to provide service to user, a webserver/standalone has been developed (https://webs.iiitd.edu.in/raghava/pep2d/).

## 1 INTRODUCTION

Over the last decade, peptides have emerged as potential therapeutic molecules against various diseases, including cancer. The peptide therapeutics has many advantages over the small molecules that include high specificity, high penetration, low production cost, ease in manufacturing and modifications [1]. In the past, various types of bioactive peptides have been reported such as hormones, toxins, neurotransmitters, antioxidants and antibiotics [1-3]. A family of peptides known as tumor homing peptides have shown enormous success in cancer treatment and diagnosis [4]. In order to manage information about peptides, a large number of peptide databases have been developed in last few years that include IEDB, CPPsite, TumorHope, HemolytiK [3-6]. Until date, about 60 peptide drugs have already released into the market and many others are under evaluation for their efficacies and toxicities in various phases of clinical trials.

Understanding the structure of bioactive peptides will enhance the understanding of peptide function as well as in developing *in silico* methods for designing peptides of desired function [7-9]. The function of a peptide can be correlated to its three dimensional (3D) or tertiary structure. In past, a large number of methods have been developed for predicting tertiary structure of proteins based on various approaches like homology, *ab initio*, threading. These methods were optimized for predicting structure of proteins not for peptides, as the hydrophobic collapse, which is a major force responsible for well-defined tertiary structure, apply well in case of proteins but not in peptides [10]. In order to overcome above limitations, methods were developed especially for predicting tertiary structure of peptides such as PEPstr [11], PEPFOLD [12-14], PEP-LOOK [15]. Prediction of secondary structure is an intermediate step in the prediction of tertiary structure and provides information about backbone. As secondary structure is the governing factor for binding characteristics of the peptide, thus an accurate prediction of secondary structure of peptides will be helpful for the scientific community. In last four decades, several methods have been developed for predicting secondary structure of proteins like Chou-Fasman, GOR, PHD, PSIPRED, Jpred [16-21].

Best of author’s knowledge no method has been developed so far for predicting secondary structure of peptides. It is possible to use protein secondary structure prediction methods for predicting secondary structure of peptides, but existing methods are trained for protein structure prediction. This raises the question whether peptide and protein adopt same secondary structure for identical segment of residues. In summary, do we need to develop separate method for predicting secondary structure of peptides or existing protein structure prediction methods are adequate. We extracted peptide structures from PDB having length less than 50 residues and compared secondary structure composition of peptides and proteins. It was observed that protein and peptide contain different percent of regular secondary structure, even identical segment adopt different secondary structure in protein and peptides. Thus, it is important to develop models for predicting secondary structure of peptides from peptide sequence. We compared the performance of newly developed models with protein secondary structure prediction methods.

## 2 METHODS

### 2.1 Datasets

#### 2.1.1 PEP2D dataset

We obtained 5778 PDB chains of peptides (2791 X-ray and 2787 NMR structures) having number of residues between 5 to 50 from ccPDB server[22]. In case structure is solved using NMR and have multiple models, we used first model of NMR. There were many peptides with unknown residues represented by letter ‘X’, we removed these peptides having more than 10% residues. We obtained 3107 unique peptides, after removing identical sequences. Our final dataset PEP2D contains 3107 unique peptide structures having 5 to 50 residues.

#### 2.1.2 PEP2D subsets

In order to understand relation between sequence and structure of peptide having length in different range, we created different subsets. First, we created a subset of smallest peptides called PEP2D5N10 that contain peptides having number of residues between 5 and 10. This subset PEP2D5N10 contains 572 peptides. Similarly, we created subset PEP2D11N20 that contains 718 peptides, having number of residues between 11 and 20. Other subsets are PEP2D21N30 (number of residues between 21 and 30) and PEP2D31N50 (number of residues between 31 and 50), containing 610 and 1207 peptides respectively.

#### 2.1.3 PEP2DNR dataset

The performance of any prediction model depends upon similarity between peptides in a dataset. In case of protein datasets, similarity level around 35% is preferred, which is not possible for peptide datasets, as it will reduce size of dataset drastically. In this study, we created a non-redundant dataset called PEP2DNR that contain 1980 peptides. This dataset only contains non-redundant peptides where no two peptides have similarity more than 80%. We used level of redundancy 80%, as previous peptide prediction also uses level of redundancy 80% [23, 24]. We used CD-HIT software [25] to make above non-redundant dataset.

#### 2.1.4 ProBlast dataset

Fundamentally, both proteins and peptides are made of 20 types of natural residues. Ideally, method developed for protein should work for peptide and vice versa. Proteins contain identical or similar peptides; question is whether isolated peptide or identical/similar peptide in protein has same structure. In order to address this question, we created a dataset called ProBlast that contains protein segment similar/identical to peptides in PEP2D. This dataset of protein segments was generated using following step; BLAST [26] search was performed for each peptide in PEP2D against protein structures in PDB [26]. The aligned region of protein segment with the peptide was extracted to make ProBlast dataset (Table S2).

### 2.2 Comparison with peptide tertiary structure prediction methods

Secondary structure of peptides can also be assigned from predicted tertiary structure of peptide. Therefore, it becomes imperative to compare our algorithm with tertiary structure prediction methods. We used PEPstr method for predicting tertiary structure of peptides in PEP2D dataset. Other popular method for peptide tertiary structure prediction (PEPFOLD) was not available as standalone and therefore, was not compared in this study. After getting tertiary structure from PEPstr, we obtained the secondary structure of peptides using DSSP software [27].

### 2.3 Peptide Secondary Structure Assignment

Secondary structure was assigned using the Dictionary of Secondary Structure of Proteins (DSSP) program. DSSP defines the distribution of secondary structure into eight different classification states [27]. We merged the H, I, G into Helix (H) state and B, E into Sheet (E) state and S, T, C into the Coil (C) state.

### 2.4 Input Features

In this study, we used binary matrix [28] and position specific scoring matrix (PSSM) as input features. In case of binary profile, each amino acid is represented by a vector of dimension 20 where one element having value one corresponding to residue type and rest of 19 elements have value zero. As there are 20 types of amino acids so each amino acid is represented by a vector of dimension 20, each element represents a residue. It means a peptide of length 8 residues will be presented by a vector of dimension 8×20. In order to use evolutionary information, we generated PSSM profiles using the HHbilts generated from multiple sequence alignments [29]. The HHbilts alignment contains up to two times more homologous sequences and results into more accurate alignments [29].

### 2.5 Classification

In this study, we applied IBK (Nearest Neighbor), Random Forest and ANN (Neural Network) machine learning techniques for prediction of peptide secondary structure. IBK and Random Forest are implemented using WEKA package [30] and ANN is implemented using monmlp (monotone multi-layer perceptron) package [31] in R statistical programming language [32].

### 2.6 Cross validation and performance measures

In this study, we performed a five-fold cross validation technique in all datasets. For assessing the performance of various models developed in this study, we computed following parameters.

#### 2.5.1 Accuracy of prediction in each state

We have calculated *Q*3*i* using equation 1, where ^*i*^ is any secondary structure element (Helix, Sheet or Coil).

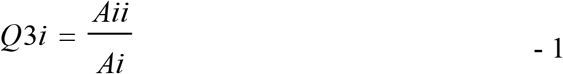

Where *Aii* is total number of correctly predicted residues in state ^*i*^, *Ai* is the total number of residues observed in state ^*i*^.

#### 2.5.2 Accuracy of prediction in all state

It is calculated by adding the number of correctly predicted residues in each state and dividing with the total number of residues in all the states (equation 2).

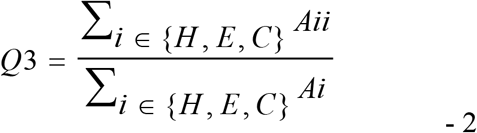

#### 2.5.3 Precision

It tells the confidence level of the prediction of each state. If the prediction accuracy is high but precision is low, this means that the prediction method is over fitted for high prediction of that state. A better prediction method should have high accuracy as well as high precision. We calculate precision as defined below in equation 3.

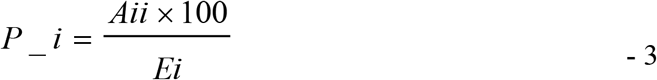

Where *Aii* is correctly predicted residues in the state ^*i*^, *Ei* is the total number of residues predicted in state ^*i*^.

#### 2.5.4 Segment Overlap Measure (SOV)

SOV represents realistic performance measure of secondary structural elements as it measures the performance at the segment level. We calculated SOV scores for each secondary structure (SOV_H, SOV_E, and SOV_C) as well as overall SOV in our predictions [33]. SOV is represented by equation 4 given below.

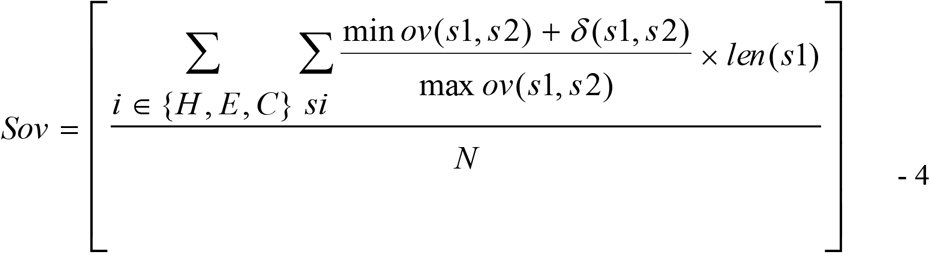

Where, *s*1 and *s*2 are segments corresponding to actual and predicted secondary structure; *len*(*s*1) corresponds to the number of residues defining the segment *s*1; min *ov*(*s*1, *s*2) corresponds to the length of overlapping *s*1 and *s*2 segments; max *ov*(*s*1, *s*2) is the maximum overlap of *s*1 and *s*2 segments for which either of the segments have a residue in state ^*i*^; *δ*(*s*1, *s*2) is a number which is defined by equation 5.

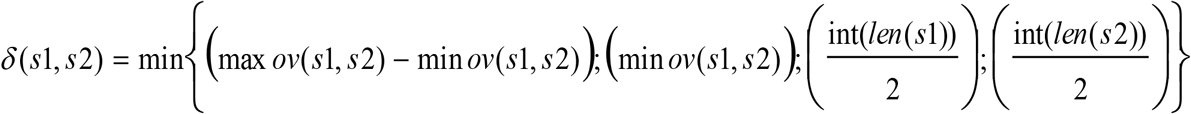

## 3 RESULTS

### 3.1 Analysis of secondary structure composition

We calculated the percent composition of Helix, Sheet and Coil for all peptides (PEP2D), similar peptide in proteins (ProBlast) and peptides having length in different range (e.g., PEP2D5N10 for length 5 to 10). The proportion of helix increases with the increase in length of the peptides, as shown in the Figure 1, percent of helix increases from 7.0 (PEP2D5N10) to 38.5% (PEP2D31N50). Similarly, coil decreases from 82.0% (PEP2D5N10) to 49.4% (PEP2D31N50). Overall, PEP2D has 33.5% helix, 10.7% sheet and 55.8% coil (Figure 1). In contrast, the ProBlast dataset has high ratio of helix 40.8%, sheet 19.9% and low coil 39.3%. This analysis indicates that similar peptides/protein segment in proteins (ProBlast) have different secondary structure composition (Table 1, Table S2).

**Table 1:** Identical sequences in peptide and protein with their PDB id and their secondary structure states.

| Amino Acid Sequence | Peptide PDB ID | Protein PDB ID | Secondary structure of peptide | Secondary structure of protein segment |
| --- | --- | --- | --- | --- |
| MSRG | 1du1A | 4aivA | CCCC | HHHC |
| YVKA | 1bfzA | 4h2kA | CCCC | HHHH |
| LDADF | 1aftA | 2zq5A | CCCCC | HHHHH |
| TPGVY | 1ab9D | 1g87A | CEEEC | CCCCE |
| CAVELR | 1b9pA | 4hacA | CCCCCC | EEEEEE |
| FVLFVL | 1b9uA | 3qyyA | HHHHHH | CEEEEE |
| PDLKAAI | 1o06A | 3visA | HHHHHHH | CCCCEEE |
| WSHPQFEK | 1rsuP | 3dp5A | CCCHHHCC | CCCCCCCC |

**Figure 1.**
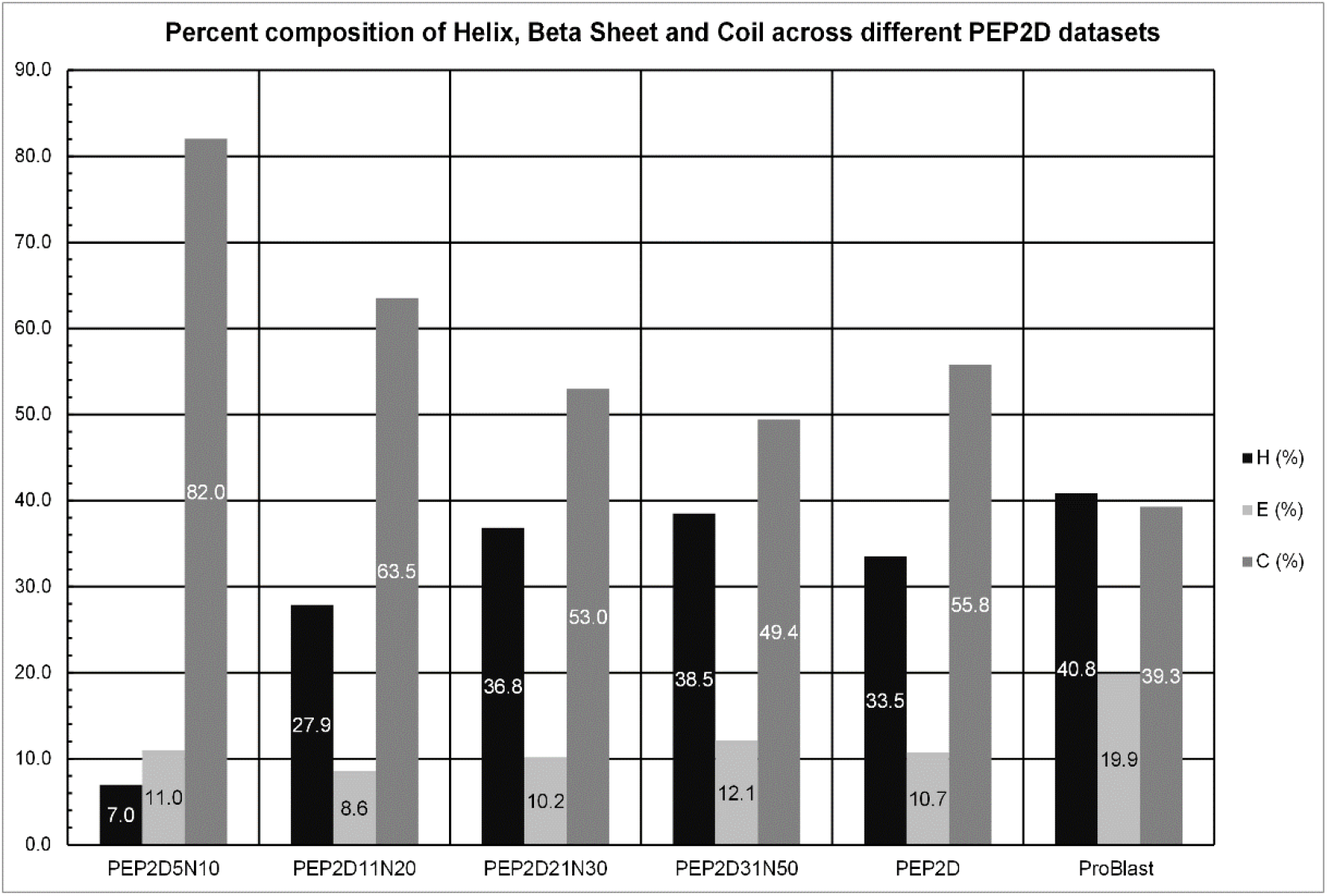
Percent composition of helix (H), sheet (E) and coil (C) in different datasets; PEP2D contain all peptides and ProBlast have peptide similar sequence in proteins. Subsets PEP2D5N10, PEP2D11N20, PEP2D21N30 and PEP2D31N50 contain peptides of length in range 5-10, 11-20, 21-30, 31-50 respectively.

### 3.2 Secondary structure of peptides in proteins

Due to their small length, peptides adopt different secondary structure as compared to protein segments that are identical to that peptide. As shown Table 1, some sequences that are identical in peptide and protein segments adopt different secondary structure. The complete list of peptides found in protein sequences are given in Supplementary file Table S2. It is because peptide in protein have different environment as their neighbor residues has an effect on their conformation. This observation emphasizes the need to develop a separate secondary structure algorithm for peptides.

### 3.3 Models based on binary profile

We developed peptide prediction models based on binary profile using different window lengths. The binary profile is represented by a vector length of W*20, where W is the length of the window (W ranges from 5-9 in this study). The best accuracy (Q3) and precision were obtained for window length 9 (WIN9). The models developed using random forest performed better than IBK and ANN based models. The random forest based model has slightly better Q3 (81.51%) as compared to IBK (79.52) and ANN based models, whereas IBK based model has better Q3E (62.63%) as compared to Q3E (47.96%) obtained by using random forest (Table 2). The precision for helix (P_H) and sheet (P_E) was also higher in case of random forest based models.

**Table 2:** The performance of models based on various techniques, developed for different window lengths using binary profile.

| Performance |  | ANN |  |  | IBK |  |  | Random Forest |  |
| --- | --- | --- | --- | --- | --- | --- | --- | --- | --- |
|  | WIN5 | WIN7 | WIN9 | WIN5 | WIN7 | WIN9 | WIN5 | WIN7 | WIN9 |
| **Q3H: Helix Accuracy** | 72.26 | 73.93 | 74.49 | 78.72 | 81.24 | 81.59 | 77.70 | 79.61 | 80.08 |
| **Q3E: Strand Accuracy** | 39.23 | 43.05 | 44.98 | 58.15 | 60.69 | 62.63 | 49.85 | 49.02 | 47.96 |
| **Q3C: Coil Accuracy** | 81.12 | 81.88 | 81.79 | 79.00 | 81.35 | 81.52 | 85.28 | 87.80 | 88.81 |
| **Q3: Overall Accuracy** | 73.66 | 75.05 | 75.40 | 76.67 | 79.10 | 79.52 | 78.94 | 80.90 | 81.51 |
| **P_H: Helix Precision** | 0.70 | 0.73 | 0.74 | 0.73 | 0.76 | 0.77 | 0.77 | 0.80 | 0.82 |
| **P_E: Strand Precision** | 0.60 | 0.57 | 0.55 | 0.62 | 0.66 | 0.65 | 0.75 | 0.81 | 0.82 |
| **P_C: Coil Precision** | 0.78 | 0.79 | 0.79 | 0.82 | 0.84 | 0.84 | 0.81 | 0.81 | 0.81 |

**Table 3:**
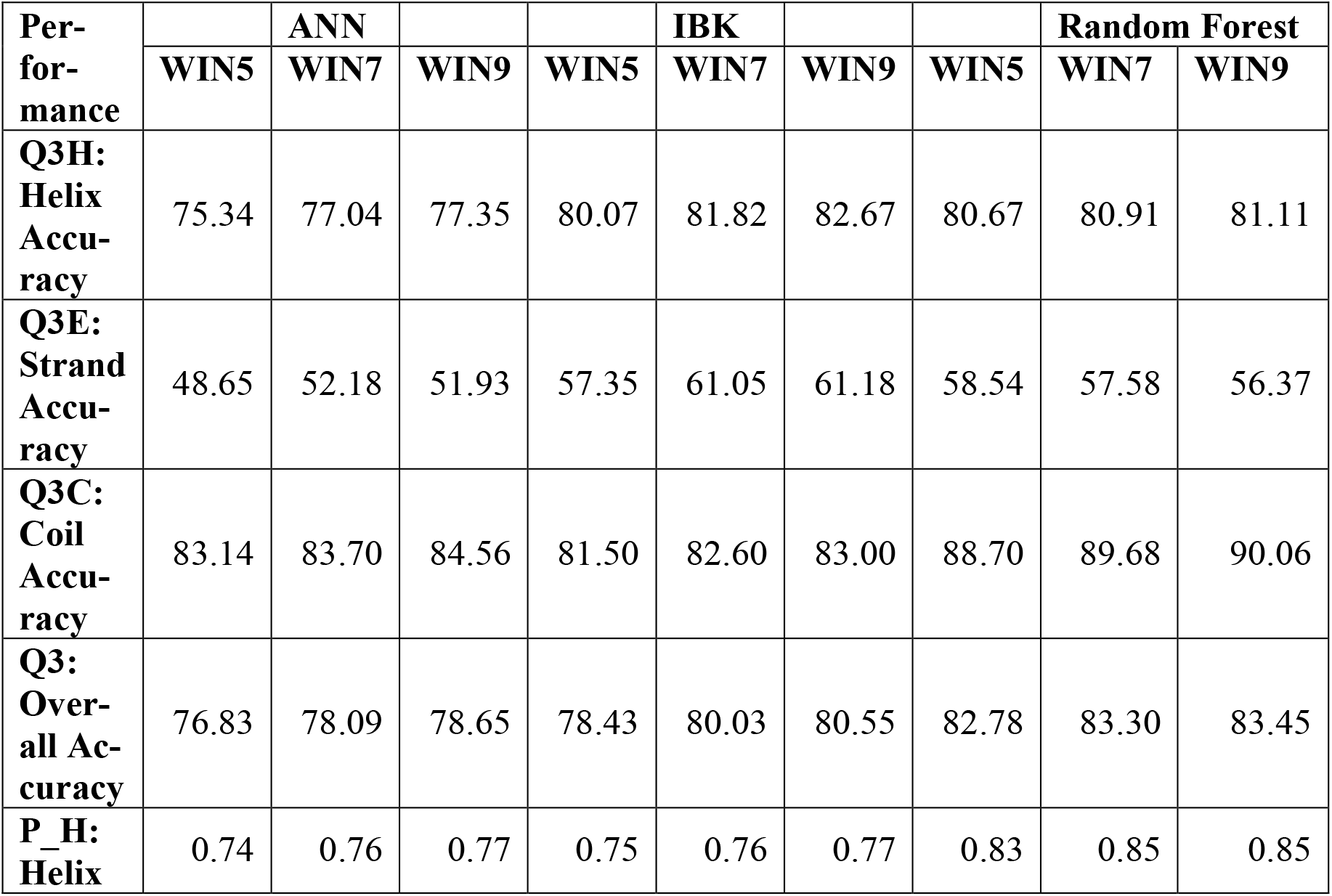
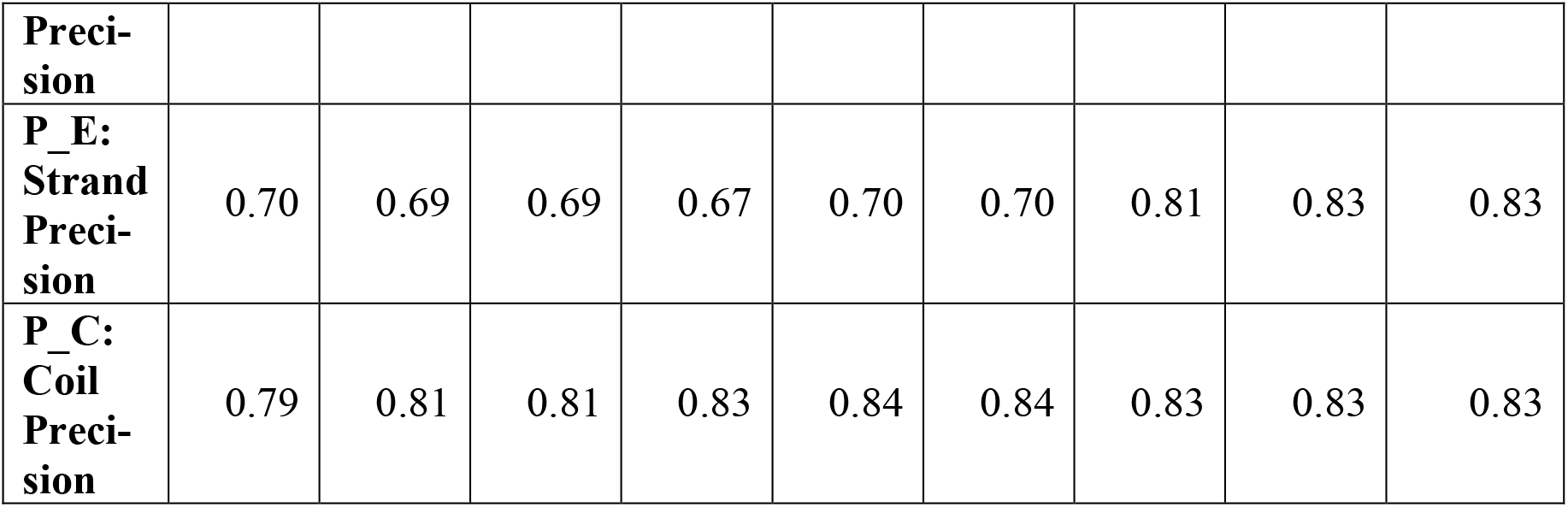
The performance of models based on various techniques, developed for different window lengths using PSSM profile.

### 3.4 PSSM profile based models

It is well known fact that evolutionary information in form of multiple sequence alignment or PSSM profile provides more information than single amino acid sequence. In past evolutionary information was used for improving performance of protein secondary structure prediction methods [18]. Our random forest based model, developed using PSSM profiles achieved overall accuracy Q3 (83.45%). Models based on random forest perform better than IBK and ANN based models for window length 9 (Table3). Again, IBK based model has higher Q3E (61.18%) as compared to random forest Q3E (56.37%). The random forest based model has considerably higher precision for helix (P_H) and sheet (P_E).

### 3.5 Balance in prediction

It was observed that in the PEP2D dataset, the content of helix and sheet was quite low as compared to coil. The more abundance of coil region resulted in a prediction model, which is biased towards high coil prediction. For obtaining a balanced prediction of helix, sheet and coil, we applied weightage to helix and sheet prediction values. The weightage values were calculated based on coil to helix and coil to sheet ratio in the dataset. The binary-based random forest model achieved maximum accuracy 80.45% (Q3) with 78.66% helix (Q3_H) and 72.03% sheet (Q3_E). The PSSM-based random forest model achieved maximum accuracy 83.48% with accuracy 80.62% for helix and 74.00% for sheet (Table 4). The performance of models improved up to 4%, when evolutionary information was used as input instead of single sequence.

**Table 4:** The performance of models developed on window 9 after applying frequency-based weightage.

| Performance | Binary |  |  | PSSM |  |  |
| --- | --- | --- | --- | --- | --- | --- |
|  | ANN | IBK | RF | ANN | IBK | RF |
| **Q3H: Helix Accuracy** | 69.55 | 76.13 | 78.66 | 76.63 | 79.77 | 80.62 |
| **Q3E: Strand Accuracy** | 63.21 | 65.32 | 72.03 | 64.01 | 65.2 | 74.00 |
| **Q3C: Coil Accuracy** | 75.60 | 83.50 | 83.13 | 80.85 | 83.78 | 87.01 |
| **Q3: Overall Accuracy** | 72.25 | 79.08 | 80.45 | 77.63 | 80.44 | 83.48 |
| **P_H: Helix Precision** | 0.78 | 0.80 | 0.83 | 0.79 | 0.81 | 0.86 |
| **P_E: Strand Precision** | 0.38 | 0.60 | 0.57 | 0.55 | 0.63 | 0.69 |
| **P_C: Coil Precision** | 0.81 | 0.82 | 0.85 | 0.82 | 0.84 | 0.85 |

### 3.6 Performance on peptides of different length

We also developed random forest model using binary and PSSM profile on different subsets of PEP2D dataset. As shown in Table 5, our binary models get overall accuracy from 78.26% to 83.45%. Similarly, PSSM based models achieved maximum accuracy between 79.04% and 84.51%. PSSM based models performed better than binary-based models in all ranges. It was also observed that difference in performance between PSSM and binary-based models improved with length of peptides. For example, for subset PEP2D5N10, difference was only 0.78% (83.45%, 84.23%) whereas for subset PEP2D31N50 difference was 2.69%. We examine PSSM profile of peptides and observed that PSSM was not properly build for small peptides up to length of 20 residues.

**Table 5:** The performance of random forest based models developed on different subsets using binary and PSSM profile.

| Performance | 5N10 |  | 11N20 |  | 21N30 |  | 31N50 |  |
| --- | --- | --- | --- | --- | --- | --- | --- | --- |
|  | Binary | PSSM | Binary | PSSM | Binary | PSSM | Binary | PSSM |
| **Q3H: Helix Accuracy** | 71.25 | 71.88 | 72.15 | 72.62 | 79.73 | 80.67 | 81.91 | 84.32 |
| **Q3E: Strand Accuracy** | 60.73 | 60.93 | 60.50 | 60.25 | 60.22 | 62.78 | 61.08 | 69.58 |
| **Q3C: Coil Accuracy** | 87.54 | 88.41 | 84.25 | 84.65 | 80.75 | 84.56 | 86.26 | 87.88 |
| **Q3: Overall Accuracy** | 83.45 | 84.23 | 78.67 | 79.04 | 78.26 | 80.88 | 81.82 | 84.51 |
| **P_H: Helix Precision** | 0.63 | 0.67 | 0.75 | 0.76 | 0.79 | 0.80 | 0.82 | 0.86 |
| **P_E: Strand Precision** | 0.46 | 0.47 | 0.51 | 0.54 | 0.59 | 0.79 | 0.77 | 0.79 |
| **P_C: Coil Precision** | 0.93 | 0.93 | 0.85 | 0.84 | 0.82 | 0.82 | 0.82 | 0.84 |

### 3.7 Comparison with PSIPRED

In order to compare performance of our method with existing methods, we evaluated performance of PSIPRED on different PEP2D datasets [20]. We used the PSIPRED (a protein secondary structure prediction method) as there is no method to predict the secondary structure of peptides. Other popular secondary structure prediction method i.e. JPred was not available as standalone. According to authors of JPred, the algorithm does not work on short peptides (<20 amino acids) as the prediction window used is 19 amino acids long and also PSSM profile obtained for these peptides is not of good quality. A benchmark performed by authors of JPred vs PSIPRED on globular proteins shows very little difference in overall accuracy [21]. In this study we have used the PSIPRED inherit in the HHsuite 2.0 [29]. This version of PSIPRED utilizes HHblits generated HMM-Profile instead of PSI-BLAST [34], which improved the accuracy of PSIPRED significantly even without retraining PSIPRED on the HHblits generated alignments as reported in HH-suite user guide. As shown in Table 6, our PSSM-based models performed better than PSIPRED on PEP2D dataset and on its subsets. Overall, on PEP2D dataset PSIPRED achieved accuracy 76.86% where as our PSSM based model achieved 83.48%. The performance of PSIPRED improves with length of peptides from 71.43% for small peptides (PEP2D5N10) to 79.80% for long peptides (PEP2D31N50). This indicates that PSIPRED is not suitable for small peptides, since it was trained on proteins. Our models perform in all ranges of peptide lengths with reasonably high accuracy. The performance of PSIPRED is particularly poor in predicting beta sheets, as accuracy is only 28.34% for small peptides (PEP2D5N10).

**Table 6:**
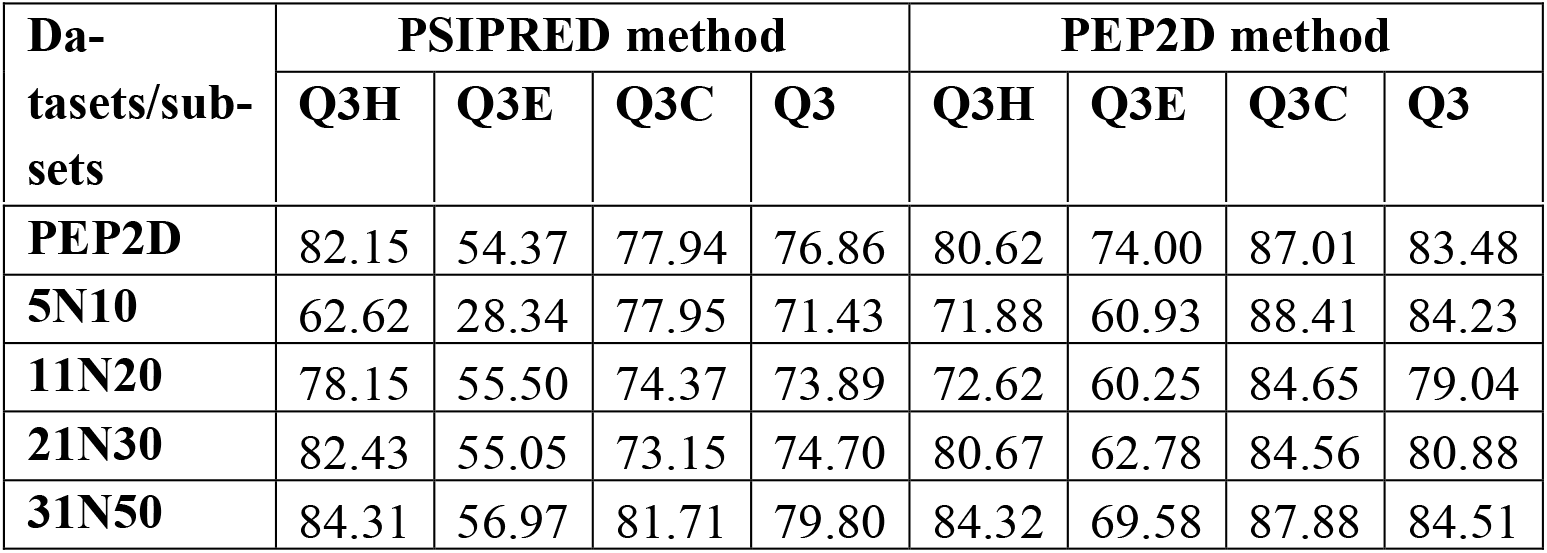
The performance of PSIPRED and PSSM based models on PEP2D dataset and its subsets.

### 3.8 Segment Overlap measure

The performance in terms of accuracy alone is not sufficient for evaluating the performance of secondary structure prediction methods. Segment Overlap (SOV) is an important measure that provides realistic procedure, which is based on evaluating the performance at the segment level. Our PSSM based model achieved SOV 76.7 with SOV_H, SOV_E and SOV_C of 75.7, 64.3 and 77.1 respectively which is much higher as compared to PSIPRED SOV 69.3. PSSM based models also perform better than PSIPRED on subsets of PEP2D particularly on small peptides (PEP2D5N10) (Table 7). As shown in Table 7, our models performed better than PSIPRED on all length of peptides.

**Table 7:** The performance of our PSSM based model and PSIPRED in terms of SOV, on dataset PEP2D and on its different subsets.

| **Da-**<br>**taset/**<br>**Sub-**<br>**sets** | **PSIPRED method** |  |  |  | **PEP2D method** |  |  |  |
| --- | --- | --- | --- | --- | --- | --- | --- | --- |
|  | **SOV_H** | **SOV_E** | **SOV_C** | **SOV** | **SOV_H** | **SOV_E** | **SOV_C** | **SOV** |
| **PEP2D** | 71.48 | 40.89 | 68.69 | 69.35 | 75.66 | 64.28 | 77.12 | 76.75 |
| **5N10** | 54.39 | 19.22 | 66.12 | 61.75 | 73.72 | 58.16 | 77.86 | 75.69 |
| **11N20** | 69.72 | 39.11 | 67.24 | 68.58 | 71.90 | 51.43 | 74.54 | 74.30 |
| **21N30** | 72.09 | 43.55 | 66.72 | 69.99 | 75.07 | 54.65 | 71.74 | 71.76 |
| **31N50** | 74.06 | 47.93 | 72.22 | 73.80 | 78.10 | 67.98 | 77.58 | 77.62 |

### 3.9 Comparison with PEPstr

We compared the performance of our models with peptide tertiary structure prediction method called PEPstr. Currently PEP-FOLD and PEPstr are state of the art methods, which are available for public use. Since PEP-FOLD, standalone was not available, so we compare our models with PEPstr. We used PEPstr for prediction of tertiary structure of all peptides in PEP2D dataset. The DSSP package was used to assign the secondary structure of peptides from their predicted tertiary structure. We achieved overall accuracy of PEPstr on PEP2D 67.09% (Q3) with accuracy 45.73% (Q3H), 2.59% (Q3E), 92.32 % (Q3C) for helix, sheet and coil respectively. The main reason of poor performance of PEPstr is that, it emphasizes on betaturn prediction over regular secondary structure. As a result, it has higher Q3C as compared to Q3H and Q3E. This comparison further strengthens the idea for a separate peptide secondary structure prediction method.

### 3.10 Performance on non-redundant dataset

The PEP2D dataset consists of unique peptides that have high level of redundancy. It is known that peptide properties can change even with a single amino acid variation. In order to understand impact of similarity between peptides in PEP2D dataset, we developed a model on non-redundant dataset (PEP2DNR). As expected, the number of peptides decreased significantly, but the performance remains significant. It was observed that only beta sheet prediction performance (Q3E) decreased to 63.5% as compared to 76.19% of PEP2D. Overall, performance of PEP2D algorithm on PEP2DNR dataset remains better than PSIPRED and lower than on PEP2D dataset (supplementary file Table S1).

## 4 IMPLEMENTATION

A user-friendly web server has been developed for predicting peptide secondary structure using our best models. The server was developed using HTML, JavaScript and PHP 5.2.9 as the front end and installed on a Red Hat Enterprise Linux 6 server environment. The server takes the FASTA sequence of single/multiple peptides and presents the results in graphical format. The server displays the query sequence, predicted secondary structure and confidence/probability score of helix, sheet and coil in graphical and text format (Figure 2). The Mutant peptide module generates all possible mutants of query peptide and predicts the secondary structure of all mutants (Figure 2). In addition, standalone version is also provided for the user, which is available at https://webs.iiitd.edu.in/raghava/pep2d/.

**Figure 2.**
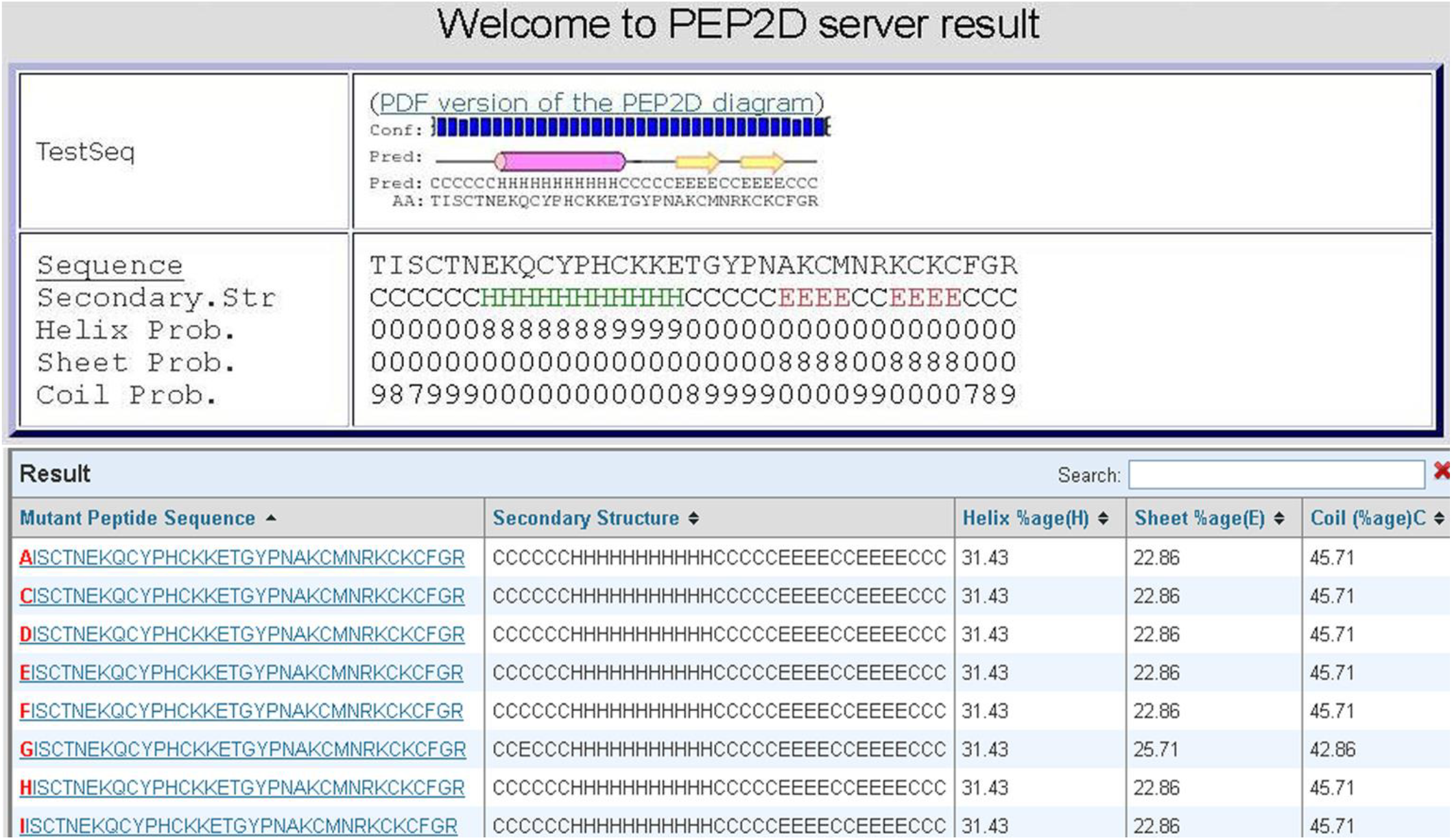
The display of result page, showing the query sequence, predicted secondary structure and probability score of helix, sheet and coil along with mutants of query sequence.

## 5 DISCUSSION

In the recent years, peptides are seen as potential therapeutic molecules due to their high specificity, high penetration, low production cost and ease in manufacturing and modifications. As secondary structure plays an important role in the function of peptides, knowing the secondary structural content of therapeutic peptides will be helpful in designing better therapeutic peptides. In past, many algorithms were developed for the prediction of secondary structure of proteins, but none for the peptides. In this study, we have developed a method PEP2D, specific for prediction of peptide secondary structure. We achieved Q3 score of 83.48% with Q3H 80.62%, Q3E 74.00% and Q3C 87.01% on PEP2D dataset. We used PSIPRED inherent in HHsuite 2.0 to predict the secondary structure of peptides of PEP2D dataset. PSIPRED was able to achieve Q3 of 76.86%, with Q3H 82.15%, Q3E 54.37% and Q3C of 77.94% on PEP2D dataset. We observed that PEP2D performs relatively better than PSIPRED particularly in the prediction of sheets. This might be due to the difference in the types/shapes of beta-sheet in peptides as compared to proteins. In peptides, the linear zig-zag conformation of the beta-sheet can be stabilized by making hydrogen bond to adjacent parallel chains of the same type. Typically, this type of beta sheet is more prominent in peptides, as bulky side chains in proteins destabilize this type of zig-zag conformation due to steric hindrance/crowding.

We created subsets of PEP2D dataset to know the effect of peptide length on accuracy. The performance improved as the length increased, as it is difficult to predict the peptides of small length due to their high irregular secondary structure content. Currently there is no algorithm to predict peptide secondary structure; we choose to compare with the state of the art method PSIPRED, for secondary structure prediction. Stating that PEP2D is better as compared to PSIPRED will not be a justifiable statement since both algorithms were trained on different types of datasets i.e. PEP2D on peptides and PSIPRED on proteins. We also compared PEP2D with peptide tertiary structure prediction method PEPstr and found that PEP2D is better in predicting secondary structure of peptides as compared to PEPstr. We hope that this method will be useful for researchers working in the field of peptide therapeutics.

## ADDITIONAL INFORMATION

Supplementary data is available online: Supplementary File.

## AUTHOR’S CONTRIBUTION

HS and SS collected and organized the data. HS and SS performed the experiments. HS and SS developed the web interface. HS and SS prepared the manuscript. GPSR conceived the idea and coordinated the project.

## ACKNOWLEDGEMENTS

Authors are thankful to Council of Scientific and Industrial Research (CSIR), Open Source Drug Discovery (OSDD) and Department of Biotechnology, Govt. of India for providing research fellowships.

## REFERENCES

1. Craik DJ, Fairlie DP, Liras S, Price D. The future of peptide-based drugs. Chem Biol Drug Des. 2013;81(1):136-47. Epub 2012/12/21. doi:10.1111/cbdd.12055. PubMed PMID:23253135.

2. Vlieghe P, Lisowski V, Martinez J, Khrestchatisky M. Synthetic therapeutic peptides: science and market. Drug Discov Today. 2010;15(1-2):40-56. Epub 2009/11/03. doi:S1359-6446(09)00370-5 [pii] 10.1016/j.drudis.2009.10.009. PubMed PMID:19879957.

3. Gautam A, Singh H, Tyagi A, Chaudhary K, Kumar R, Kapoor P, et al. CPPsite: a curated database of cell penetrating peptides. Database (Oxford). 2012;2012:bas015. Epub 2012/03/10. doi:bas015 [pii] 10.1093/database/bas015. PubMed PMID:22403286; PubMed Central PMCID:PMC3296953.

4. Kapoor P, Singh H, Gautam A, Chaudhary K, Kumar R, Raghava GP. TumorHoPe: a database of tumor homing peptides. PLoS One. 2012;7(4):e35187. Epub 2012/04/24. doi:10.1371/journal.pone.0035187 PONE-D-12-00452 [pii]. PubMed PMID:22523575; PubMed Central PMCID:PMC3327652.

5. Vita R, Zarebski L, Greenbaum JA, Emami H, Hoof I, Salimi N, et al. The immune epitope database 2.0. Nucleic Acids Res. 2010;38(Database issue):D854-62. Epub 2009/11/13. doi:gkp1004 [pii] 10.1093/nar/gkp1004. PubMed PMID:19906713; PubMed Central PMCID:PMC2808938.

6. Gautam A, Chaudhary K, Singh S, Joshi A, Anand P, Tuknait A, et al. Hemolytik: a database of experimentally determined hemolytic and non-hemolytic peptides. Nucleic Acids Res. 2014;42(Database issue):D444-9. Epub 2013/11/01. doi:gkt1008 [pii] 10.1093/nar/gkt1008. PubMed PMID:24174543.

7. Sharma A, Kapoor P, Gautam A, Chaudhary K, Kumar R, Chauhan JS, et al. Computational approach for designing tumor homing peptides. Sci Rep. 2013;3:1607. Epub 2013/04/06. doi:srep01607 [pii] 10.1038/srep01607. PubMed PMID:23558316; PubMed Central PMCID:PMC3617442.

8. Gautam A, Chaudhary K, Kumar R, Sharma A, Kapoor P, Tyagi A, et al. In silico approaches for designing highly effective cell penetrating peptides. J Transl Med. 2013;11:74. Epub 2013/03/23. doi:1479-5876-11-74 [pii] 10.1186/1479-5876-11-74. PubMed PMID:23517638; PubMed Central PMCID:PMC3615965.

9. Lata S, Mishra NK, Raghava GP. AntiBP2: improved version of antibacterial peptide prediction. BMC Bioinformatics. 2010;11 Suppl 1:S19. Epub 2010/03/05. doi:1471-2105-11-S1-S19 [pii] 10.1186/1471-2105-11-S1-S19. PubMed PMID:20122190; PubMed Central PMCID:PMC3009489.

10. Bradley P, Chivian D, Meiler J, Misura KM, Rohl CA, Schief WR, et al. Rosetta predictions in CASP5: successes, failures, and prospects for complete automation. Proteins. 2003;53 Suppl 6:457-68. Epub 2003/10/28. doi:10.1002/prot.10552. PubMed PMID:14579334.

11. Kaur H, Garg A, Raghava GP. PEPstr: a de novo method for tertiary structure prediction of small bioactive peptides. Protein Pept Lett. 2007;14(7):626-31. Epub 2007/09/28. PubMed PMID:17897087.

12. Thevenet P, Shen Y, Maupetit J, Guyon F, Derreumaux P, Tuffery P. PEP-FOLD: an updated de novo structure prediction server for both linear and disulfide bonded cyclic peptides. Nucleic Acids Res. 2012;40(Web Server issue):W288-93. Epub 2012/05/15. doi:gks419 [pii] 10.1093/nar/gks419. PubMed PMID:22581768; PubMed Central PMCID:PMC3394260.

13. Maupetit J, Derreumaux P, Tuffery P. PEP-FOLD: an online resource for de novo peptide structure prediction. Nucleic Acids Res. 2009;37(Web Server issue):W498-503. Epub 2009/05/13. doi:gkp323 [pii] 10.1093/nar/gkp323. PubMed PMID:19433514; PubMed Central PMCID:PMC2703897.

14. Lamiable A, Thevenet P, Rey J, Vavrusa M, Derreumaux P, Tuffery P. PEP-FOLD3: faster de novo structure prediction for linear peptides in solution and in complex. Nucleic Acids Res. 2016;44(W1):W449-54. doi:10.1093/nar/gkw329. PubMed PMID:27131374; PubMed Central PMCID:PMCPMC4987898.

15. Thomas A, Deshayes S, Decaffmeyer M, Van Eyck MH, Charloteaux BB, Brasseur R. PepLook: an innovative in silico tool for determination of structure, polymorphism and stability of peptides. Adv Exp Med Biol. 2009;611:459-60. Epub 2009/04/30. PubMed PMID:19400265.

16. Chou PY, Fasman GD. Prediction of protein conformation. Biochemistry. 1974;13(2):222-45. Epub 1974/01/15. PubMed PMID:4358940.

17. Sen TZ, Jernigan RL, Garnier J, Kloczkowski A. GOR V server for protein secondary structure prediction. Bioinformatics. 2005;21(11):2787-8. Epub 2005/03/31. doi:bti408 [pii] 10.1093/bioinformatics/bti408. PubMed PMID:15797907; PubMed Central PMCID:PMC2553678.

18. Rost B, Sander C, Schneider R. PHD--an automatic mail server for protein secondary structure prediction. Comput Appl Biosci. 1994;10(1):53-60. Epub 1994/02/01. PubMed PMID:8193956.

19. Rost B, Sander C. Prediction of protein secondary structure at better than 70% accuracy. J Mol Biol. 1993;232(2):584-99. Epub 1993/07/20. doi:S0022-2836(83)71413-0 [pii] 10.1006/jmbi.1993.1413. PubMed PMID:8345525.

20. McGuffin LJ, Bryson K, Jones DT. The PSIPRED protein structure prediction server. Bioinformatics. 2000;16(4):404-5. Epub 2000/06/27. PubMed PMID:10869041.

21. Cole C, Barber JD, Barton GJ. The Jpred 3 secondary structure prediction server. Nucleic Acids Res. 2008;36(Web Server issue):W197-201. Epub 2008/05/09. doi:gkn238 [pii]10.1093/nar/gkn238. PubMed PMID:18463136; PubMed Central PMCID:PMC2447793.

22. Singh H, Chauhan JS, Gromiha MM, Raghava GP. ccPDB: compilation and creation of data sets from Protein Data Bank. Nucleic Acids Res. 2012;40(Database issue):D486-9. Epub 2011/12/06. doi:gkr1150 [pii]10.1093/nar/gkr1150 [doi]. PubMed PMID:22139939; PubMed Central PMCID:PMC3245168.

23. Singh H, Ansari HR, Raghava GP. Improved method for linear B-cell epitope prediction using antigen’s primary sequence. PLoS One. 2013;8(5):e62216. Epub 2013/05/15. doi:10.1371/journal.pone.0062216 PONE-D-13-06566 [pii]. PubMed PMID:23667458; PubMed Central PMCID:PMC3646881.

24. El-Manzalawy Y, Dobbs D, Honavar V. Predicting flexible length linear B-cell epitopes. Comput Syst Bioinformatics Conf. 2008;7:121-32. Epub 2008/01/01. PubMed PMID:19642274; PubMed Central PMCID:PMC3400678.

25. Li W, Godzik A. Cd-hit: a fast program for clustering and comparing large sets of protein or nucleotide sequences. Bioinformatics. 2006;22(13):1658-9. Epub 2006/05/30. doi:btl158 [pii] 10.1093/bioinformatics/btl158. PubMed PMID:16731699.

26. Altschul SF, Gish W, Miller W, Myers EW, Lipman DJ. Basic local alignment search tool. J Mol Biol. 1990;215(3):403-10. Epub 1990/10/05. doi:10.1016/S0022-2836(05)80360-2S0022-2836(05)80360-2 [pii]. PubMed PMID:2231712.

27. Kabsch W, Sander C. Dictionary of protein secondary structure: pattern recognition of hydrogen-bonded and geometrical features. Biopolymers. 1983;22(12):2577-637. Epub 1983/12/01. doi:10.1002/bip.360221211 [doi]. PubMed PMID:6667333.

28. Agarwal S, Mishra NK, Singh H, Raghava GP. Identification of mannose interacting residues using local composition. PLoS One. 2011;6(9):e24039. Epub 2011/09/21. doi:10.1371/journal.pone.0024039 PONE-D-11-05490 [pii]. PubMed PMID:21931639; PubMed Central PMCID:PMC3172211.

29. Remmert M, Biegert A, Hauser A, Soding J. HHblits: lightning-fast iterative protein sequence searching by HMM-HMM alignment. Nat Methods. 2012;9(2):173-5. Epub 2011/12/27. doi:nmeth.1818 [pii] 10.1038/nmeth.1818 [doi]. PubMed PMID:22198341.

30. Mark Hall EF, Geoffrey Holmes, Bernhard Pfahringer, Peter Reutemann, Ian H. Witten. The WEKA Data Mining Software: An Update;. SIGKDD Explorations. 2009;11(1).

31. Cannon AJ. Monotone multi-layer perceptron neural network. R package version 1.1.2. 2012.

32. Team RDC. R: A language and environment for statistical computing. R Foundation for Statistical Computing, Vienna, Austria. ISBN 3-900051-07-0 2011.

33. Zemla A, Venclovas C, Fidelis K, Rost B. A modified definition of Sov, a segment-based measure for protein secondary structure prediction assessment. Proteins. 1999;34(2):220-3. Epub 1999/02/18. doi:10.1002/(SICI)1097-0134(19990201)34:2<220::AID-PROT7>3.0.CO;2-K [pii]. PubMed PMID:10022357.

34. Altschul SF, Madden TL, Schaffer AA, Zhang J, Zhang Z, Miller W, et al. Gapped BLAST and PSI-BLAST: a new generation of protein database search programs. Nucleic Acids Res. 1997;25(17):3389-402. Epub 1997/09/01. doi:gka562 [pii]. PubMed PMID:9254694; PubMed Central PMCID:PMC146917.

